# InTraSeq: A Multimodal Assay that Uncovers New Single-Cell Biology and Regulatory Mechanisms

**DOI:** 10.1101/2024.09.19.613947

**Authors:** Majd M. Ariss, Linglin Huang, Xiaokai Ding, Shivani Sheth, Tyler Levy, Jeremy Fisher, Jean Loebelenz, Keith Arlotta, Karen Dixon, Roberto Polakiewicz, Vijay K. Kuchroo, Sean A. Beausoleil

**Affiliations:** Cell Signaling Technology, Inc., Danvers, MA 01923, USA; The Gene Lay Institute of Immunology and Inflammation, Brigham and Women’s Hospital, Massachusetts General Hospital and Harvard Medical School, Boston, MA 02115, USA; Klarman Cell Observatory, Broad Institute of MIT and Harvard, Cambridge, MA 02142, USA; Department of Biomedicine, University of Basel, 4031 Basel, Switzerland

**Author notes:** Co-first authors (Majd M. Ariss, Linglin Huang, Xiaokai Ding). Correspondence: Sean A. Beausoleil Vijay K. Kuchroo.

**Keywords:** InTraSeq, single-cell, intracellular protein, post-translational modification, signaling pathway

## Abstract

Single-cell RNA sequencing (scRNA-seq) has revolutionized cell biology by enabling the profiling of transcriptomes at a single-cell resolution, leading to important discoveries that have advanced our understanding of cellular and tissue heterogeneity, developmental trajectories, and disease progression. Despite these important advances, scRNA-seq is limited to measuring the transcriptome providing a partial view of cellular function. To address this limitation, multimodal scRNA-seq assays have emerged, allowing for the simultaneous measurement of RNA expression and protein. Intracellular Transcriptomic and Protein Sequencing (InTraSeq), a novel multimodal scRNA-seq technology described here, enables the concurrent measurement of mRNA, surface markers, cytoplasmic proteins, and nuclear proteins within individual cells through oligo-barcoded antibodies. This method offers a comprehensive approach to studying cellular function by combining RNA and protein profiling from the same sample and utilizing a relatively simple protocol. The InTraSeq method enables researchers to expand their view of critical intracellular protein expression including post-translational modifications (PTMs) and transcription factors, allowing for the identification of novel cellular subtypes and states that may be obscured by RNA-based analyses alone. This is particularly valuable in understanding the heterogeneity of cell populations and identifying distinct functional states. In this report, we used InTraSeq to characterize the complex cellular states and regulatory mechanisms during Th17 cell differentiation. We simultaneously profiled RNA and protein expression in over 85,000 cells, capturing transcriptional changes, changes in protein expression and the dynamics of signaling pathways at a high resolution. Our results revealed novel insights into Th17 cell differentiation, including the identification of key regulatory factors and their target genes. By simultaneously measuring mRNA, extra and intra-cellular proteins, signaling proteins, and PTMs, InTraSeq offers a comprehensive understanding of cellular processes and enables the identification of novel regulatory mechanisms.

## Introduction

Single-cell RNA sequencing (scRNA-seq) has emerged as a transformative tool in cell biology, enabling the profiling of transcriptomes at single-cell resolution^1–5^. This technology has revolutionized our understanding of cellular heterogeneity, allowing for the identification of novel cell types, tracking of cell differentiation trajectories, and elucidation of the molecular groundwork of development, disease, and immunity^6–10^. While scRNA-seq offers valuable insights into cellular gene expression^11,12^, achieving a holistic understanding of cellular function necessitates measuring proteins, as they play a substantial role in driving many cellular processes that dictate behavior and phenotype^13^. Additionally, understanding the interplay between RNA expression, protein expression, and post translational modifications (PTMs) is crucial for dissecting complex signal transduction pathways and characterizing the molecular mechanisms behind disease development^14–16^.

Due to the importance of measuring proteins along with mRNA, recent advances in single-cell multimodal assays have emerged as a powerful strategy to tackle the limitations of scRNA-seq. These assays enable the simultaneous measurement of multiple cellular features including RNA expression, protein levels, chromatin accessibility, and epigenetic modifications^17,18^. However, integrating multiple molecular modalities from the same cell presents technical challenges. For instance, capturing both RNA and intracellular protein in single cell datasets often compromises RNA integrity^19,20^ and requires joint embedding with other snRNA-seq datasets to obtain an acceptable single-cell RNA readout^20^. Some multimodal approaches require bridging oligos and are not optimized for cytoplasmic protein readouts^21^. Other methods resort to using nuclei instead of whole cells as their input, resulting in the loss of cytoplasmic proteins and the majority of information from signaling pathways^17,20^.

Additionally, recent studies bypass some of these challenges by constructing separate single-cell libraries for each modality followed by computationally bridging the separate libraries together^18,21^. However, this approach presents additional challenges due to the need for complex computational methods and algorithms and increases the amount of lab benchwork for each additional single cell library all of which introduce opportunities for misinterpretation of results.

The computational integration makes the implicit assumption that the two libraries are exact replicates with identical cell compositions, and that the association between chromatin accessibility and gene expression is strong enough to dominate over cell-to-cell heterogeneity. The specificity of these assumptions raises concerns about the quality of the integration and the interpretability of the data.

To address all these limitations and challenges, we introduce InTraSeq (Intracellular protein and Transcriptomic Sequencing), a novel technology with a simple experimental workflow. InTraSeq allows for the simultaneous measurement of RNA, surface markers, cytoplasmic proteins, and nuclear proteins inclusive of PTMs within individual cells, enabling insights into altered protein activity, stability, and expression levels. This capability enables researchers to expand their single cell experiments to include important information about protein expression and post-translational modification typical of signaling molecules at single-cell resolution. Importantly, InTraSeq maintains high-quality measurement of mRNA consistent with current scRNA-seq methods within individual cells without compromising the quality of either data modality. This streamlined experimental workflow eliminates the need to create multiple single-cell multimodal libraries. In addition, the InTraSeq provides us with the ability to integrate activation of intracellular signaling components to the downstream transcriptional activation within the same cell. In this report we demonstrate InTraSeq’s utility to advance our understanding of single-cell biology as exemplified by its application to Th17 T-cell differentiation.

## Results

### InTraSeq generates high-quality single-cell RNA and protein profiles in the same cells

The InTraSeq 3’ assay concurrently profiles the transcriptome and proteome in the same cells at single-cell resolution. InTraSeq uses a buffer kit which processes the cells for a total of one hour of benchwork spread out over three days (Figure 1A, Supplementary Protocol 1). First, the cells are fixed with Methanol overnight. At this point the cells can be stored for up to 7 days at -20C°. After fixation, the cells are incubated with the “scBlock” buffer (Cat #82906) and then immunostained with antibody oligo conjugates for 16 hours. On the third day, the cells were washed three time with a Wash Buffer before being subjected to a scRNA-seq using the10X

**Figure 1.**
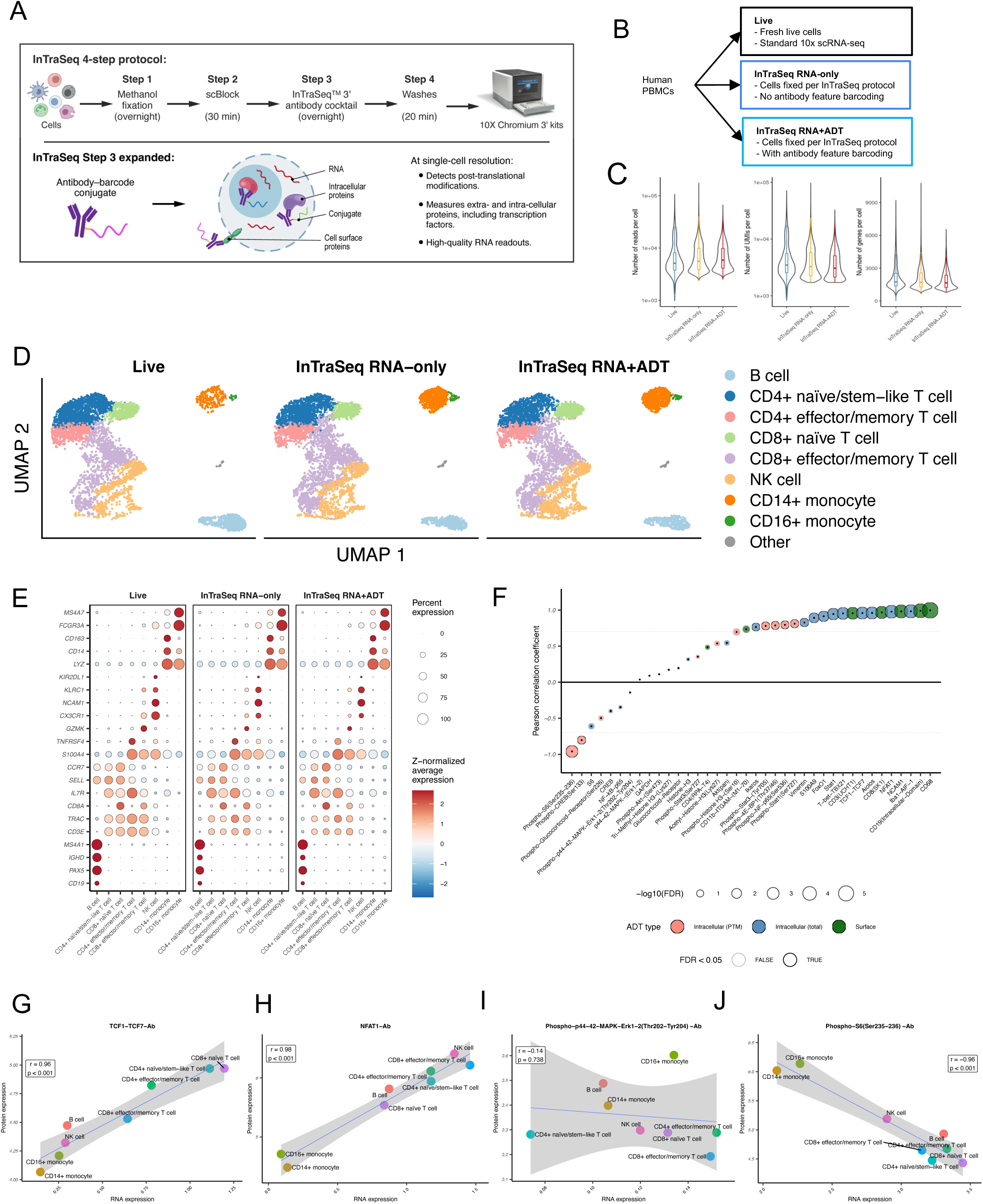
InTraSeq generates high-quality single-cell RNA and protein profiles in the same cells. A. InTraSeq workflow. B. Experiment design of the benchmarking analysis. C. Distribution of summary statistics of the gene expression assay comparing the Live, InTraSeq RNA-only and InTraSeq RNA+ADT samples. To exclude impacts of the differences in the sequencing depth and the number of cells captured, only 7,000 cells with largest total number of reads were included in the comparison, and the samples were down-sampled to match the median number of reads per cell in the sample with lowest coverage (Methods). D. UMAP plots depicting the cell types identified in the Live, InTraSeq RNA-only and InTraSeq RNA+ADT samples. UMAP coordinates were computed based on RNA expression data after integration. Colors indicate the cell types. E. RNA expression of cluster marker genes in each sample. Size of dots reflects percent of cells with non-zero UMIs, and the color represents average expression of genes that were z-normalized across cell types in each sample. F. Correlation between average RNA and protein expression across cell types in the InTraSeq RNA+ADT data. Size and border color of dots reflect the statistical significance of the Pearson correlation coefficient. Fill color of the dots shows the protein types. G-J. Average RNA (x-axis) vs. protein (y-axis) expression across cell types for TCF1 (G), NFAT (H), phospho MAPK-Erk1-2 (I) and phospho S6 (J). Pearson correlation coefficients and corresponding unadjusted p-values were reported for each RNA-protein pair.

Genomics 3’ assay (Cat#CG000317). In order to effectively enable antibody capture without interfering with mRNA capture, we leveraged the 10XGenomics Feature Barcoding technology in the oligo design (see methods).

To demonstrate whether InTraSeq can generate high-quality single-cell mRNA profiles, the transcriptomes in human peripheral blood mononuclear cells (PBMCs) from the same donor were benchmarked by performing a standard 10x Genomics Chromium Single Cell 3’ v3.1 experiment in the following three conditions: 1-live PBMCs (Live), 2-PBMCs that underwent the InTraSeq 3’ protocol without the addition of antibody feature barcodes (InTraSeq RNA-only), and 3-PBMCs that underwent the InTraSeq 3’ protocol with antibody feature barcodes (InTraSeq RNA+ADT) (Figure 1B). After matching on the mean reads per cell for the top 7,000 cells with highest total UMI counts, the InTraSeq RNA-only and InTraSeq RNA+ADT samples displayed similar total UMIs per cell and number of genes per cell compared to the Live PBMCs sample (Figure 1C, Supplementary table 2, Methods).

Deeper analysis of the RNA profiles shows that RNA expression levels were tightly correlated between the InTraSeq samples and the live control (Supplementary Figure 1A). The three samples were integrated using the standard Harmony integration pipeline with default parameters^22^ (Supplementary Figure 1B). Unsupervised clustering analysis showed the three datasets have similar cell type compositions, and the cell type relative frequencies are consistent with the previously published experiments^23^ (Figure 1D, Supplementary Figure 1C, Table S3). Similar expression patterns of top cell type marker genes across datasets were also observed, such as *CD19*, *CD3E*, *CD8A*, *SELL*, *NCAM1*, *CD14*, and *FCGR3A*, further validating high fidelity in mRNA expression between InTraSeq and 10x scRNA-seq data (Figure 1E, Methods). Replication of this experiment yielded consistent results across three biological replicates (Supplementary Figure 1D-F).

Furthermore, the RNA and protein expression in the InTraSeq RNA+ADT data were compared at the cell type level. As expected, significantly positive correlations between RNA and protein abundance were observed for the many of the targets, particularly the surface proteins such as CD68, CD19, NCAM1 and CD8, and some intracellular proteins such as AIF1, NFAT1, and TCF7 (Figure 1F-H). However, the Pearson correlation coefficient rapidly decreased for many intracellular proteins such as Glucocorticoid Receptor (GR), MAPK-ERK1-2, CREB1 (|r| < 0.5, FDR ≥ 0.05). Additionally, 10 out of 14 of the antibodies against post-translational modifications (PTMs) tended to exhibit minimal (FDR ≥ 0.05) or even negative correlations with the expression of their encoding genes (Figure 1F, I, J). This was not surprising since PTMs are regulated at the protein levels and cannot be inferred from RNA expression alone. This highlights the unique advantage of InTraSeq, which enables the measurement of PTMs at the single-cell level, particularly when high quality PTM specific antibodies are available providing novel insights that are inaccessible through scRNA sequencing alone.

### InTraSeq captures extra- and intra-cellular protein signals to distinguish cell types and reveal cell states

To assess the quality and robustness of the InTraSeq protein signal, a deeper analysis of the InTraSeq RNA+ADT sample was performed. As demonstrated in Figure 1, the high-quality mRNA expression data itself allowed the identification of different cell types (Figure 2A). While many of the protein and mRNA expression patterns were consistent in feature plots and observed in the same cluster of cells in the UMAP, the protein data exhibited less sparsity compared to the RNA data (Figure 1F, Figure 2B, Supplementary Figure 1B-C, and Table S4). To validate the InTraSeq protein signal of S100A9, NFAT1, and AIOLOS, an independent flow cytometry analysis was conducted in a separate PBMC sample. The flow results were similar to the InTraSeq dataset showing S100A9 to be expressed in monocytes, NFAT1 being predominantly observed in T and NK cells with low expression in B cells, and AIOLOS being predominantly expressed in B cells then NK and T cells (Figure 2C). The cross-validation using flow cytometry demonstrates the reliability of the InTraSeq single-cell protein data.

**Figure 2.**
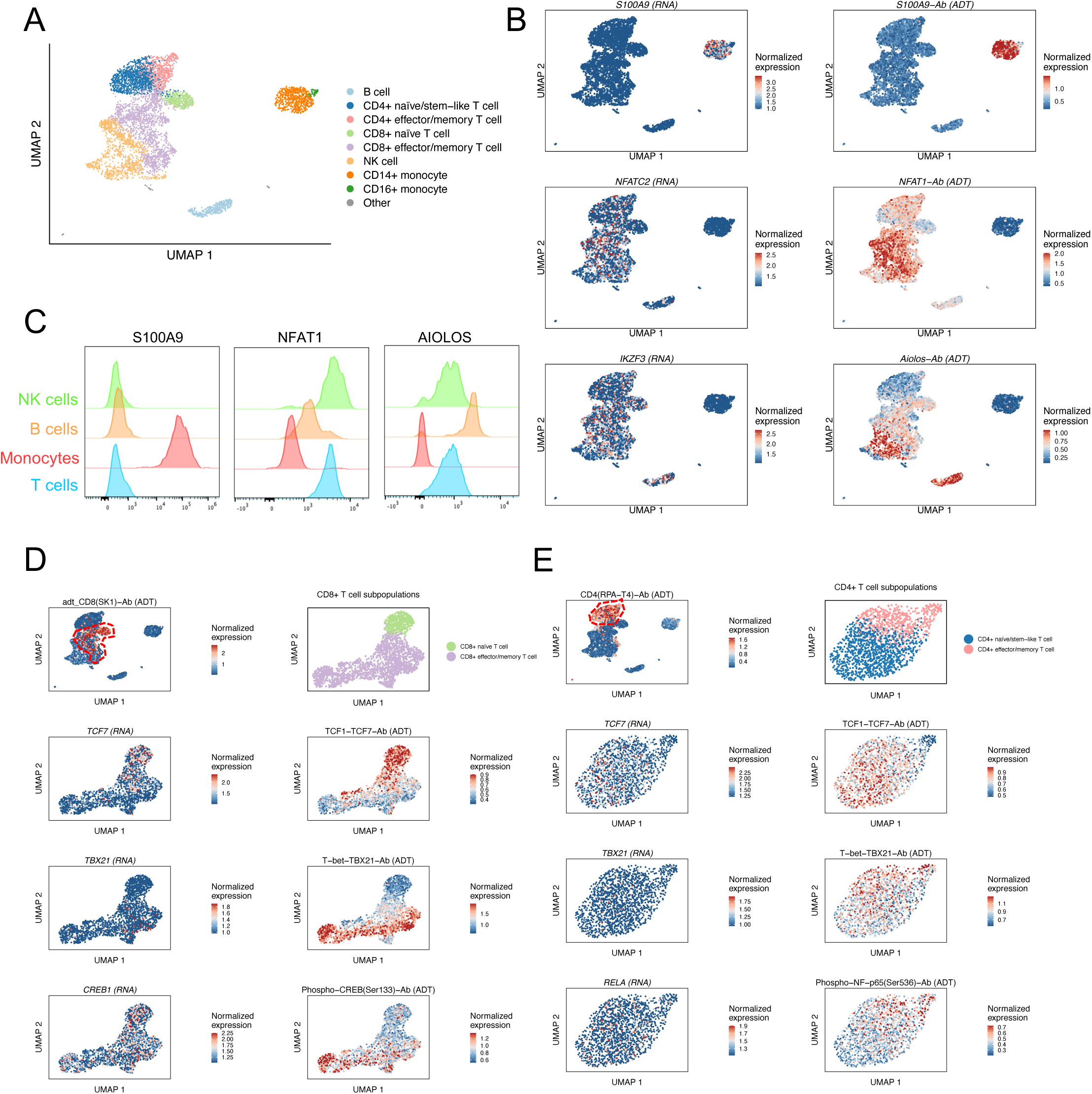
InTraSeq captures extra- and intra-cellular protein signals to distinguish cell types and unveil intricate cell states. A. UMAP plot depicting the cell types in the InTraSeq RNA+ADT sample. UMAP coordinates were computed based on RNA expression data of the sample. Colors indicate the cell types. B. UMAP plots depicting the expression of genes of interest (RNAs) and their corresponding proteins (ADTs). The RNA data were log-normalized within every cell, and the ADT data were central-log-ratio-normalized across all cells. C. Flow cytometry validation of the expression patterns across cell types for proteins of interest. D. UMAP plots showing the CD8+ T cell states and the expression of genes and proteins of interest. The top-left panel showed the CD8 protein expression and the location of CD8+ T cell clusters in the whole InTraSeq RNA+ADT data. The top-right panel showed the two CD8+ T cell subpopulations. The UMAP coordinates for all panels other than top-left were computed based on RNA data of the two CD8+ T cell clusters. E. UMAP plots showing the CD4+ T cell states and the expression of genes and proteins of interest. The top-left panel showed the CD4 protein expression and the location of CD4+ T cell clusters in the whole InTraSeq RNA+ADT data. The top-right panel showed the two CD4+ T cell subpopulations. The UMAP coordinates for all panels other than top-left were computed based on RNA data of the two CD4+ T cell clusters.

To determine whether InTraSeq’s intracellular protein readout can reveal novel biological insights, mRNA and protein expression were analyzed in different cellular states of one cell type. A CD8+ T cell focused analysis was performed to compare two CD8+ cellular states: a “naïve” population characterized by high *CCR7*, and *SELL* transcript expression and an “effector/memory” population defined by high *S100A4* and *GZMK* transcript levels ^24,25^ (Figure 1E). TCF1, a known protein marker of T cell stemness, and TBX21, a known transcription factor associated with effector/memory T cells exhibited distinct protein expression patterns in the “naïve/stem” and “effector/memory” cell populations respectively on the UMAP, aligning with previous studies^26,27^ (Figure 2D). Additionally, phosphorylated CREB1 (Ser133) was enriched in effector/memory CD8+ T cells, consistent with its role in T cell activation^28^ (Figure 2D). In contrast, RNA expression of *TCF7* and *TBX21* was sparse and lacked clear differentiation across CD8+ T cell states (Supplementary Figure 2D), reinforcing the utility of both protein and PTM readouts to effectively define differential cell states.

A similar analysis of CD4+ T cells revealed analogous protein expression patterns for TCF1 and TBX21 in line with previous studies^26,29^, with corresponding RNA data showing sparser signals (Figure 2E and Supplementary Figure 2E). Furthermore, phosphorylated NF-κB (Ser536), linked to T cell activation^30^, was enriched in effector CD4+ T cells at the protein level but not at the RNA level (Figure 2E and Supplementary Figure 2E).

These findings demonstrate InTraSeq’s ability to detect robust proteomic signals, enabling deeper exploration of cellular heterogeneity and identification of cell states, particularly for genes with low RNA expression or proteins regulated by post-translational modifications.

### InTraSeq reveals transcriptomic and proteomic dynamics during Th17 cell differentiation

In addition to characterizing cell states, InTraSeq’s simultaneous measurement of intracellular proteins, post translational modifications, and transcripts facilitates the investigation of signaling pathway activation and subsequent transcriptional changes upon stimulation at a single-cell resolution. To demonstrate this, the Th17 cell differentiation was studied using InTraSeq.

Previous studies have shown that the Th17 cell differentiation pathway activation occurs within minutes of stimulation^31^, while the transcriptional profile changes initiate within hours and evolves over days^32^. To capture the dynamics of both the signaling pathways and the gene expression, naïve CD4+ T cells from mouse spleens and lymph nodes were cultured under Th0 differentiation conditions (anti-CD3 + anti-CD28), non-pathogenic Th17 differentiation conditions (npTh17; anti-CD3 + anti-CD28 + IL-6 + TGFβ), pathogenic Th17 differentiation conditions (pTh17; anti-CD3 + anti-CD28 + IL-6 + IL-1β + IL-23) or PMA and Ionomycin (PMA/IO) stimulation (anti-CD3 + anti-CD28 + PMA + Ionomycin). Cells were collected at 0 minutes (naïve cells), 10 minutes, 45 minutes, 6 hours and 24 hours post-stimulations (Figure 3A). The PMA/IO stimulation condition served as a positive control since they potently activate multiple signaling cascades bypassing the need for TCR engagement and costimulatory signals, with data collected at 10 and 45 minutes.

**Figure 3.**
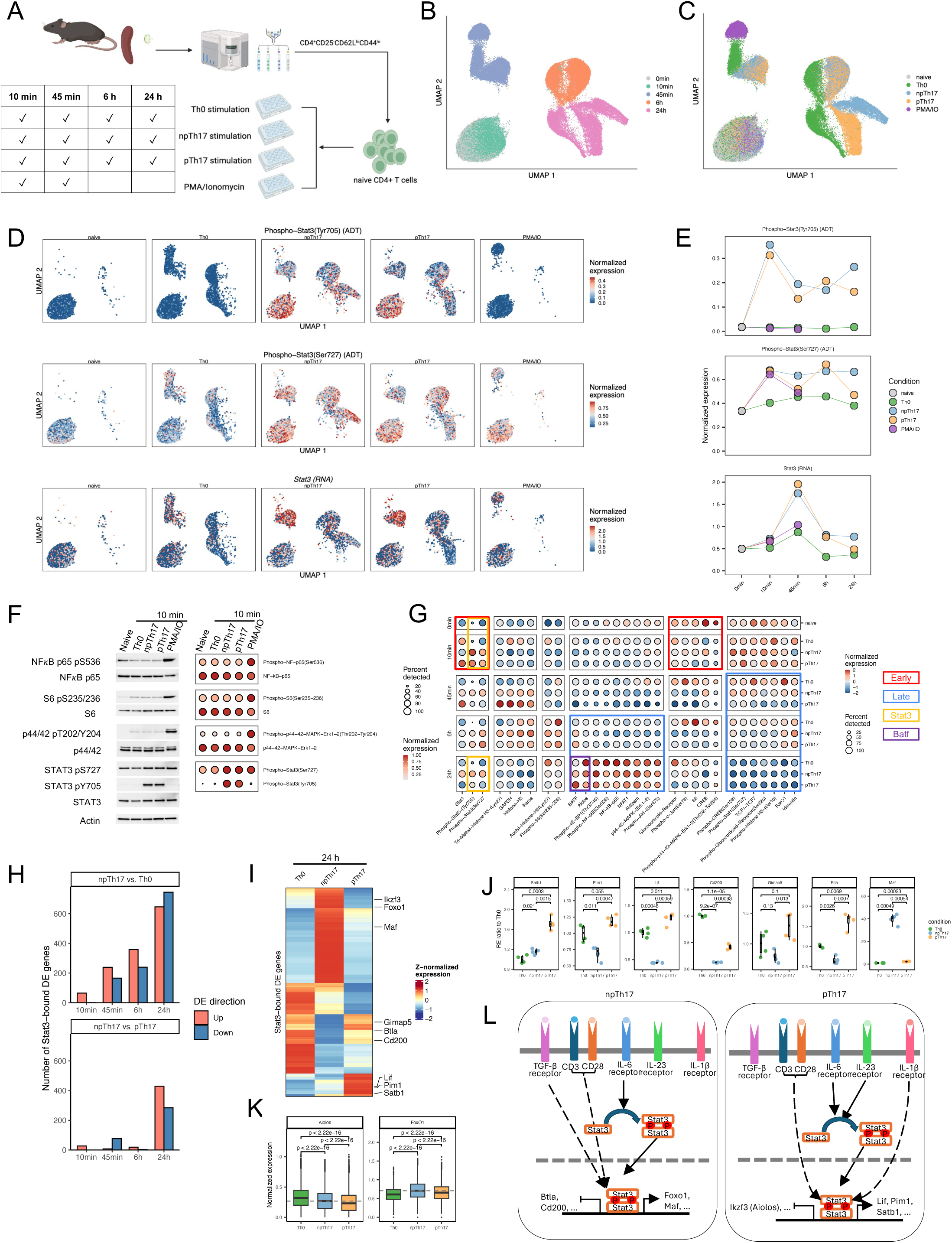
InTraSeq captures the proteomics signals and transcriptional changes during Th17 cell differentiation. A. Schematic of experiment design. B. UMAP plot showing the cells of different time points. UMAP coordinates were computed using RNA data. C. UMAP plot showing the cells of different stimulations. The UMAP coordinates are the same as Figure 3B. D. UMAP plots showing the phospho Stat3 (Ty705) protein, the phospho Stat3 (Ser727) protein, and Stat3 RNA expression in each stimulation. The UMAP coordinates are the same as Figure 3B. E. Average expression of phospho Stat3 (Ty705) protein, the phospho Stat3 (Ser727) protein, and Stat3 RNA in every condition over time. F. A comparison of the Western blot validation results (left) and InTraSeq data (right) for selected proteins showed substantial change at 10 minutes compared to naïve CD4+ T cells. G. Dot plot depicting the expression of all proteins at every time and stimulation combination except PMA/IO treatment. Proteins were ordered and clustered using hierarchical clustering. H. Number of Stat3-bound genes that are significantly up- or down-regulated in npTh17 compared to Th0 (top) or pTh17 (bottom) at each time point. I. Expression of Stat3 direct targets of interest at the 24-hour time point. The color indicates the z-normalized average expression. J. qPCR validations of selected Stat3 direct targets of interest. Y-axis represents the ratio of a sample’s relative expression (RE) compared to the mean RE of the Th0 group. K. Expression of Stat3 direct targets of interest that were included in the InTraSeq antibody panel. L. Schematic of non-pathogenic and pathogenic Th17 differentiation pathways according to the Stat3-focused analysis.

The InTraSeq Th17 dataset profiled RNA and protein expression in 83,772cells across 16 samples (4,243 - 6,811 cells per sample, Table S5). Unsupervised clustering analysis based on mRNA data revealed distinct clusters from different timepoints, with transcriptional differences between Th0 and Th17 differentiation conditions emerging at 45 minutes (Figure 3B, Supplementary Figure 3A-B). The pTh17 and npTh17 conditions were transcriptionally distinct only at 24 hours, indicating delayed cytokine effects on the pathogenic state driven by IL-1β and IL-23, and the non-pathogenic states driven by TGFβ (Figure 3C, Supplementary Figure 3A-B).

Additionally, the 10-minute post-stimulation samples did not cluster separately from the naïve CD4+ T cells since the differences in the transcription profiles were insufficient to drive distinct clusters at this early timepoint (Figure 3B, Supplementary Figure 3A-B). In contrast, protein-based single-cell clustering demonstrated pronounced clustering difference at 10 minutes between naïve and differentiated cells, highlighting InTraSeq’s ability to quantify early proteomic changes not observable using RNA alone (Supplementary Figure 3C-F). The contribution of the abundance of PTM-specific antibodies in InTraSeq enabled the detection and visualization of early acute proteomic changes, revealing novel insights into complex cellular dynamics prior to the activation of the transcriptional machinery.

Previous studies have emphasized the critical role of Stat3 in Th17 differentiation. This process is initiated by IL-6 binding to its receptor, IL-6R, which subsequently activates JAK1 and JAK2 kinases. These activated kinases then phosphorylate Stat3, triggering its dimerization and translocation to the nucleus where it induces the transcription of genes involved in Th17 cell differentiation^33,34^. The time course InTraSeq data enabled observation of dynamic changes in phosphorylated Stat3 (pStat3-Tyr705 and pStat3-Ser727), and *Stat3* mRNA relative to naïve and Th0 samples (Figure 3D-E). Consistent with previous findings, pStat3-Tyr705 levels significantly increased and peaked in npTh17 and pTh17 conditions as early as 10 minutes after stimulation, persisting at a slightly lower level until 24 hours. While pStat3-Ser727 also increased at 10 minutes, this change was observed in PMA/IO stimulation, suggesting involvement of mechanisms beyond the IL-6 signaling pathway, such as TCR activation. *Stat3* RNA increased at 45 minutes, with highest levels in npTh17 and pTh17 conditions, followed by a sharp decline at 6 hours. The distinct temporal patterns of pStat3 protein and RNA levels support its known positive autoregulatory role, wherein Stat3 activation in CD4+ T cells drives its own expression by binding to the *Stat3* gene promoter^35^.

To validate the pStat3 signal, Western blot analysis was performed on naïve T cells and stimulated Th0, npTh17, pTh17, and PMA/IO cells collected 10 minutes post-stimulation. Consistent with InTraSeq protein data, pStat3-Tyr705 upregulation was observed exclusively in Th17 samples (Figure 3F). Similarly, pNF-κB-Ser536, pS6-Ser235/236, and pMAPK-ERK1/2-Thr202/204 were upregulated in PMA/IO samples by both Western blot and InTraSeq analysis(Figure 3F). Cross-validation with Western blot supports the robustness of the InTraSeq single-cell phosphoprotein data, and demonstrates the ability to simultaneously measure mRNA, protein, and PTM expression in a single experiment.

Furthermore, additional InTraSeq protein readouts in the Th17 experiment revealed diverse patterns across conditions and time points (Figure 3G). Notably, activation and repression modules were identified based on protein expression dynamics. For example, FoxO1 and Tcf1 transcription factor protein levels decreased at the 45 minutes timepoint, with a substantial drop in the Th17 stimulation conditions relative to Th0 (Figure 3G, Supplementary Figure 3G).

Conversely, BATF and Aiolos both increased after 24 hours. However, BATF showed a larger increase in pTh17 compared to npTh17 and Th0 at 24 hours, while Aiolos had the highest increase in the Th0 sample at the same time point (Figure 3G, Supplementary Figure 3H-I).

To elucidate regulatory mechanisms in Th17 cells, Stat3 and BATF were selected as example targets (Figure 3H-K and Supplementary Figure 3K-N). Given pStat3-Tyr705 upregulation in Th17 differentiation, direct Stat3 target genes were identified using published Stat3 ChIP-seq data from npTh17 cells to determine Stat3-bound gene signatures^36^ (Methods). The number of Stat3-bound genes that are differentially expressed in npTh17 vs. Th0 gradually increased at each timepoint, peaking at 24h, with 626 upregulated genes and 745 downregulated genes (FDR < 0.05, fold change > 1.5), demonstrating the impact of Stat3 in the Th17 differentiation system (Fig 3H).

Additionally, the InTraSeq experiment also showed decreased pStat3 levels in pTh17 relative to npTh17 at 24 hours (Figure 3E). To further dissect this, a deeper analysis was performed to determine whether the Stat3-bound genes are differentially expressed between the npTh17 and pTh17 samples. The results show a striking upregulation of 429 and downregulation of 284 Stat3-bound genes exclusively at the 24h timepoint (Fig. 3H).

To identify Stat3 direct targets that play a role in Th17 differentiation and associated with Th17 pathogenicity, Stat3-bound npTh17 vs. Th0 differential genes (Fig.3H, top) were intersected with Stat3-bound npTh17 vs. pTh17 differential genes (Figure 3H, bottom).

These Stat3 direct targets of interest were differentially regulated by pStat3 signaling perturbations in non-pathogenic and pathogenic Th17 conditions (Figure 3I). For instance, robust transcriptional changes occurred with *Satb1*, *Pim1*, *Lif*, *Cd200*, *Gimap5*, and *Btla* being upregulated in pTh17 compared to npTh17, and *Maf* being downregulated, with the data being further validated using qPCR (Figure 3I-J, Figure 3L).

To further elucidate the pStat3 signaling mechanism in the pathogenicity of Th17 cells, two direct targets of Stat3, *Ikzf3* (encoding gene for the Aiolos protein) and *Foxo1* (encoding gene for the FoxO1 protein) were found to be significantly downregulated at the transcript level in pTh17 relative to npTh17 at 24 hours (Figure 3I). Given that antibodies against Aiolos and FoxO1 are in the InTraSeq panel, the protein expression levels of these two targets were compared across conditions at the 24-hour timepoint. Consistent with the mRNA data, the InTraSeq protein levels for Aiolos and FoxO1 were also downregulated in pTh17 (Figure 3K), supporting the differential expression of these direct Stat3-bound genes. This finding underscores the ability of InTraSeq to deconvolute subtle differences in signaling pathways, such as the downregulation of Aiolos and FoxO1 in pTh17 compared to npTh17, which is driven by altered Stat3 signaling and supported by both mRNA and protein data (Figure 3L).

A similar analysis was performed focusing on BATF, given its late activation and differential expression between pTh17 and npTh17 conditions (Figure 3G). Using publicly available ChIP-seq data^36^, BATF-bound target genes were identified and a subset of the genes were found to be differentially expressed in npTh17 vs. Th0 and/or pTh17 at the 24-hour time point (Supplementary Figure 3M-N; Methods).

### InTraSeq allowed identification of mitotic cells using phosphorylated Histone H3 (Ser10) abundance

Cell cycle regulation is crucial for immune cell differentiation and expansion. Performing a default cell cycle scoring based analysis on the single cell mRNA data identified proliferating cells (S, G2, and M phase) (Methods) primarily at the 24 hours timepoint in the Th17 experiment, consistent with expected cell cycle synchronization following prolonged cell culturing (Figure 4A). To refine cell cycle phase distribution, cell cycle signature scores were recalculated respectively for every sample at 24 hours, revealing 7.8%-15.9% of cells in S phase, 61.9%-64.1% of cells in G1 phase, and 22.2%-28.7% of cells in G2M phase (Figure 4B-C).

**Figure 4.**
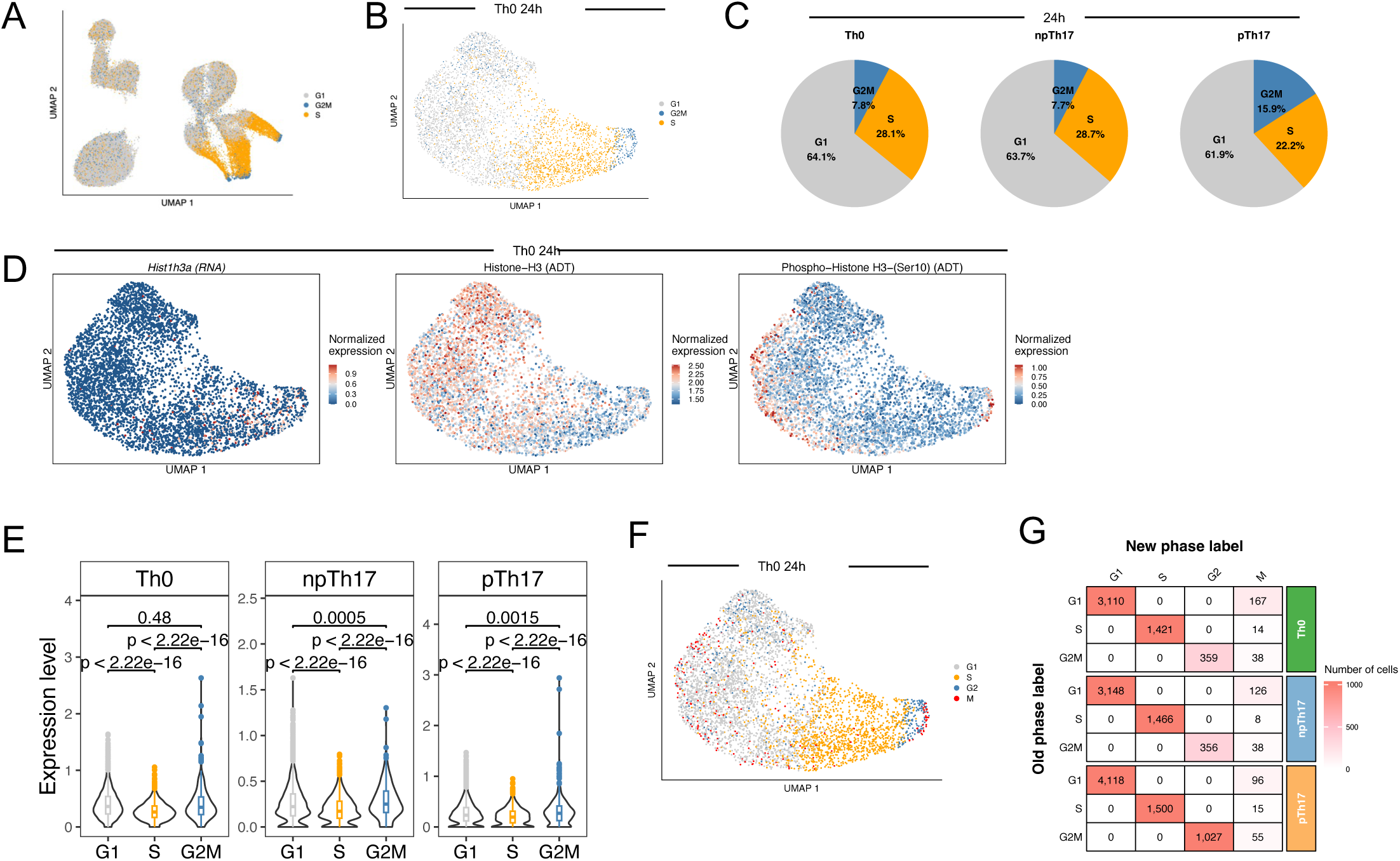
InTraSeq identifies mitotic cells with post-translational modification of Histone H3. A. UMAP plot showing the cell phase labels for all samples based on RNA data. UMAP coordinates were the same as Figure 3B. B. UMAP plot showing the cell phase labels re-annotated for the Th0 stimulation sample at 24 hours. Re-annotation and UMAP coordinates were computed based on the RNA data of the Th0 at 24-hour data. C. Pie chart showing the proportion of cells belong to every cell cycle phases after re-annotation. D. UMAP plots showing the expression of Hist1h3a gene (RNA), total Histone H3 protein, and pH3-Ser10 protein in Th0 at 24 hours. E. Distribution of pH3-Ser10 protein levels in cells at different cell cycle phases at 24 hours. F. UMAP plot showing the new cell cycle phase annotation using additional information from pH3-Ser10 protein levels in Th0 at 24 hours. G. A comparison of the old and new cell cycle phase labels in each sample at 24 hours. The numbers are the number of cells belong to the category.

Histone H3 modifications are linked to dynamic chromatin and cell cycle progression^37^. Specifically, phosphorylation of histone H3 on serine 10 (pH3-Ser10) marks chromosome condensation during mitosis^38^. To investigate cell cycle dynamics, RNA expression of Histone H3 encoding gene (*Hist1h3a*), total H3 protein, pH3-Ser10, and the pH3-Ser10 to total H3 protein ratio were examined (Figure 4D, Supplementary Figure 4A-B). While *Hist1h3a* expression significantly correlated with S and G2M gene signature scores (Pearson 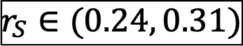 and 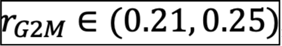, respectively; all p-values < 2.2×10^-16^), total H3 protein levels showed negative correlations (Pearson 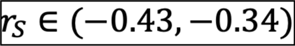 and 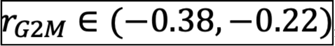, respectively; all p-values < 2.2×10^-16^). pH3-Ser10 (and its ratio to H3) was enriched in a subset of the G2M and G1 phase cells (Figure 4D-E, Supplementary Figure S4A-C). This discrepancy between RNA, protein, and PTM data suggested limitations in assessing cell cycle states using mRNA alone. To address this, cells with high pH3-Ser10 levels were annotated as mitosis (M) phase cells (Methods). This approach identifies 4.3%, 3.3% and 2.4% of cells as M phase cells in Th0, npTh17 and pTh17 samples respectively at the 24-hour timepoint (Figure 4F-G, Supplementary Figure 4D-E), demonstrating InTraSeq’s sensitivity in capturing transient cell cycle phases. The relatively low percentage of cells in the M phase (4.3%-7.8%) is consistent with the shorter duration of mitosis compared to interphase (G1, S, and G2 phases)^39^. This finding underscores the importance of InTraSeq for accurately capturing transient cell states that may be overlooked by mRNA-based analyses alone.

## Discussion

This study demonstrates the effectiveness of InTraSeq for comprehensive RNA and protein profiling in both human and mouse cells, offering valuable insights into cellular biology. The PBMC dataset showcased accurate and robust mRNA and protein measurements, further validated by flow cytometry. The use of PBMC cell types as internal controls ensured the specificity and sensitivity of antibody-based protein detection in InTraSeq.

InTraSeq’s ability to measure transcription factors represents a significant advancement over traditional single-cell mRNA sequencing methods. Transcription factor (TF) transcripts are notoriously difficult to detect in single-cell datasets due to their low abundance and rapid turnover^40^. InTraSeq addresses this limitation by enabling the measurement of key TFs such as TCF1, NFAT1, TBX21, and Aiolos. This capability provides valuable insights into cellular differentiation and function, as TFs play pivotal roles in regulating gene expression and cellular identity.

In circumstances where mRNA and protein expression patterns diverge, InTraSeq excels in identifying and characterizing cellular states that were often overlooked in traditional mRNA-based analyses. For instance, InTraSeq robustly labeled CD4/CD8 states, enabling the characterization of subpopulations within one defined cell type (Figure 2D-E). This capability is particularly valuable in cancer biology, where malignant cells may exhibit diverse states not readily characterized by mRNA expression alone.

This study applied InTraSeq to profile RNA and protein expression during CD4+ Th17 cell differentiation, capturing signaling activities through intracellular protein analysis, particularly PTMs, prior to significant transcriptional changes. Using Stat3 as an example, we demonstrated InTraSeq’s ability to identify signaling and transcriptional regulatory pathways, focusing on early-stage (within 24 hours) non-pathogenic and pathogenic Th17 differentiation. Despite the established role of Stat3 in Th17 differentiation, novel *Stat3* target genes potentially influencing Th17 pathogenicity were identified.

Furthermore, InTraSeq’s ability to identify novel regulatory mechanisms is exemplified by the discovery of a potential role for BATF in Th17 differentiation. BATF is a transcription factor known to be involved in T cell differentiation, but its specific role in Th17 cells is not well understood. By analyzing protein expression dynamics and identifying BATF target genes, InTraSeq revealed a novel role for BATF in regulating specific gene expression programs associated with Th17 pathogenicity.

InTraSeq’s ability to measure acute proteomic changes at the post-translational modification level, even within short timeframes where RNA expression remains relatively unchanged, underscores its value in studying rapid cellular perturbations. For example, the Th17 cell differentiation experiment demonstrated InTraSeq’s ability to quantify phospho-Stat3-Tyr705, phospho-Stat3-Ser727, phospho-p44/42-Thr202/Tyr204, phospho-S6-Ser235/236, and phospho-NFκB-Ser536 at 10 minutes, a time point where RNA changes were minimal (Figure 3E, 3G). This highlights the importance of InTraSeq for detecting early-stage signaling pathway activities and cellular responses that precede transcriptional alterations.

While the study provides valuable insights, it has limitations. The Th17 differentiation data were derived from pooled cells without cell hashing or repeated experiments, affecting statistical analysis. To address this, findings were validated using Western blot and qPCR. Future studies could benefit from incorporating cell hashing and biological replicates to enhance statistical rigor. Additionally, expanding the antibody panel in InTraSeq would enable a more comprehensive analysis of Th17 cell states, such as the inclusion of markers specific to pathogenic and non-pathogenic Th17 differentiation.

In the Stat3 and BATF downstream target analysis, the successful use of publicly available Stat3 and BATF ChIP-seq data supports the reliability of InTraSeq findings. While the data timepoints do not perfectly align with the InTraSeq experimental conditions, the observed correlations between ChIP targets and InTraSeq data validate the approach. Future studies could generate in-house ChIP-seq data for each timepoint for a more precise alignment with InTraSeq experiments.

This study contributes to the understanding of signaling transduction and gene transcription regulation during T cell activation and cytokine signaling. InTraSeq’s ability to measure protein levels, post-translational modifications of signaling proteins, and transcriptomes at the single-cell level provides novel insights into the interactions between these molecular entities.

---

**Supplementary Figure 1.**
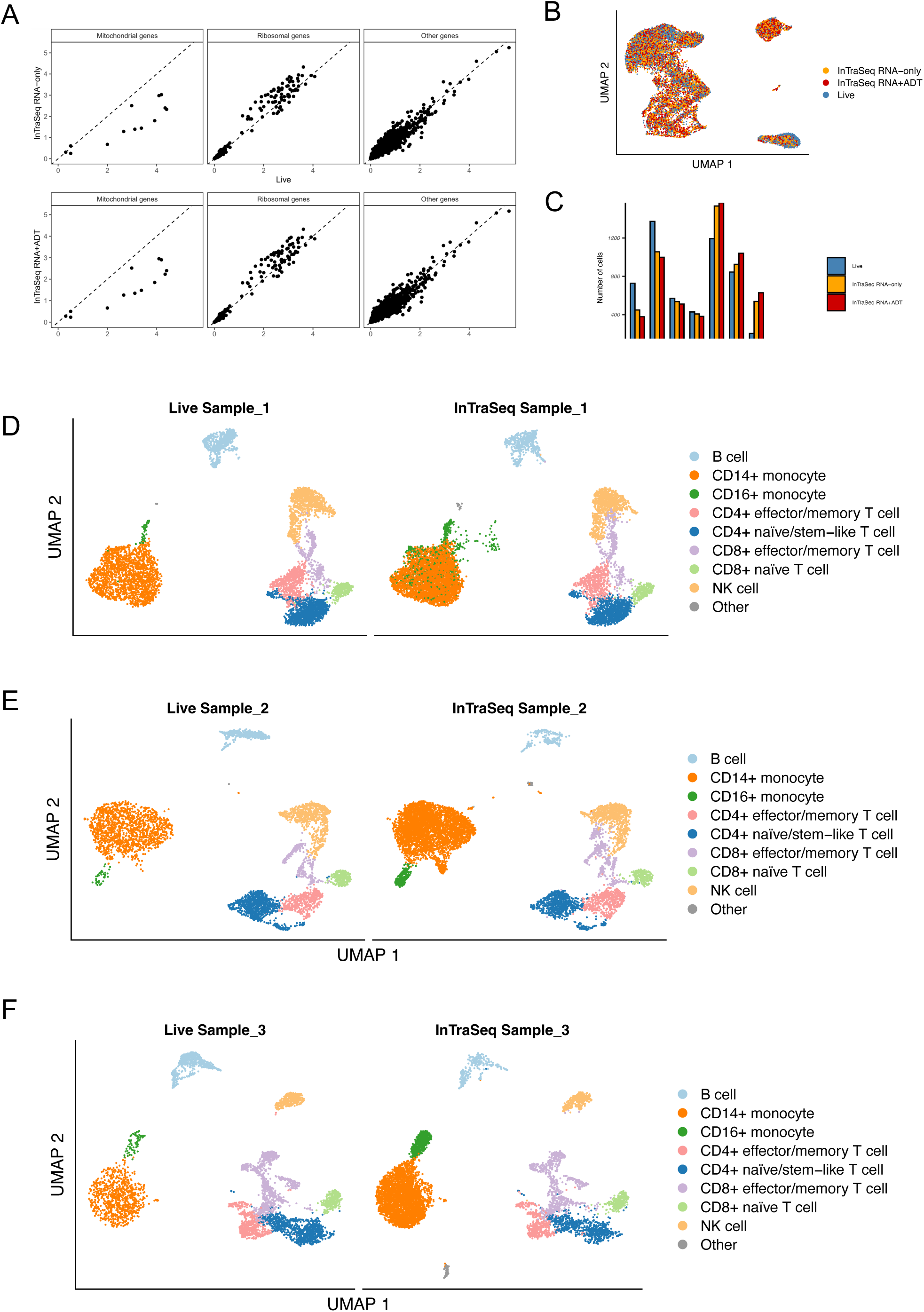
A. Scatter plots comparing the average log-normalized expression of genes in Live vs. InTraSeq RNA-only (top) or vs. InTraSeq RNA+ADT (bottom). Genes were stratified into three types, and the gene type labels were obtained from the GENCODE vM23 annotation. B. UMAP plot showing the mixing of three samples after Harmony integration (Methods). C. Bar plot comparing the number of cells belong to different cell types in the three samples. D. UMAP plots showing the cell types in Live and InTraSeq data from an independent experiment on a different PBMC sample (Sample_1). E. UMAP plots showing the cell types in Live and InTraSeq data from an independent experiment on a different PBMC sample (Sample_2). F. UMAP plots showing the cell types in Live and InTraSeq data from an independent experiment on a different PBMC sample (Sample_3).

**Supplementary Figure 2.**
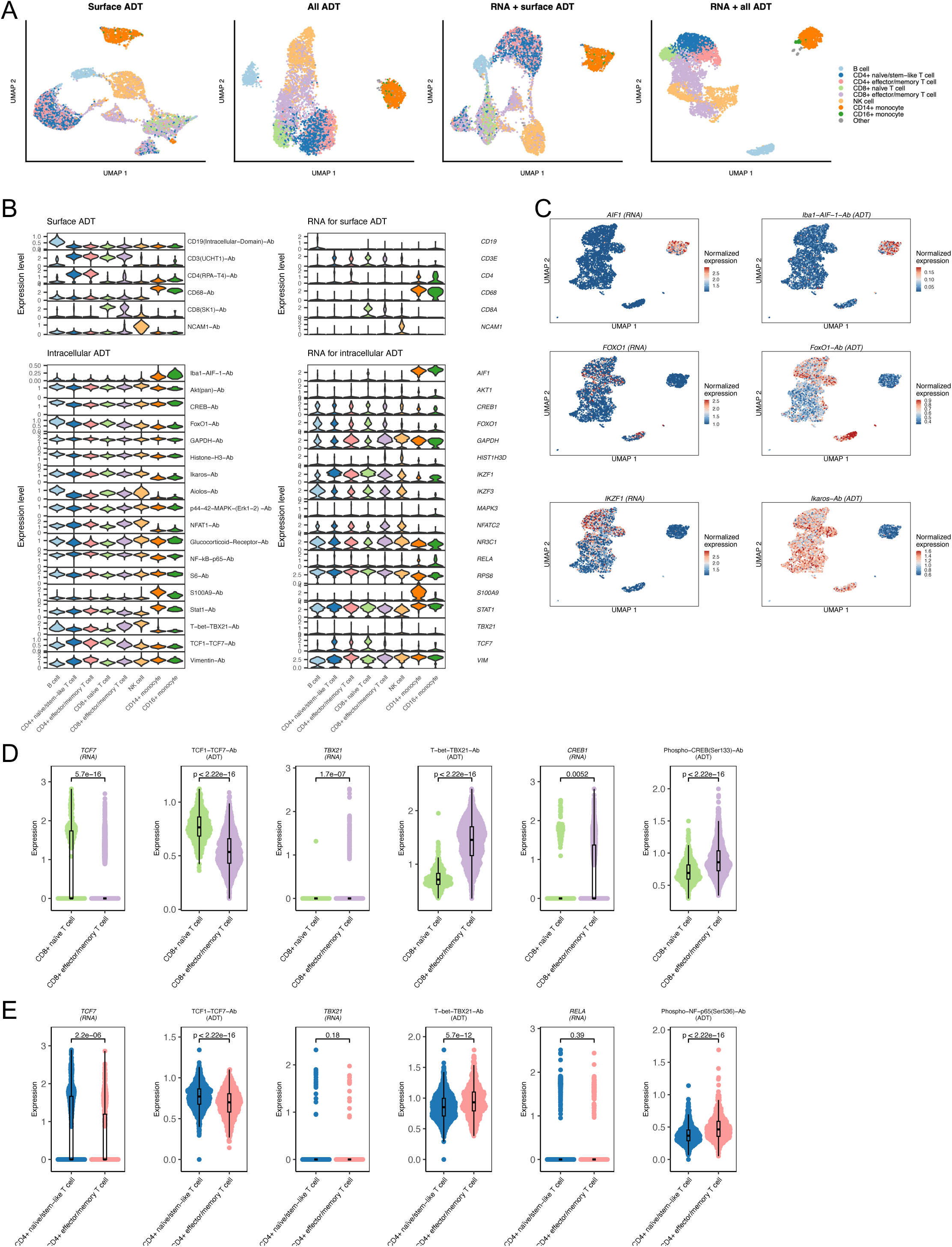
A. UMAP plots for the cells from the PBMC InTraSeq RNA+ADT sample. UMAP coordinates were computed based on surface protein data only (left 1), all protein data (left 2), RNA and surface protein data (left 3), and RNA and all protein data (right 1). B. Stacked violin plots showing the expression patterns of the cell surface proteins (top left) and their corresponding RNAs (top right), and the intracellular total proteins (bottom left) and their corresponding RNAs (bottom right) across cell types in the InTraSeq RNA+ADT sample. C. UMAP plots depicting the expression of genes of interest (RNAs) and their corresponding proteins (ADTs). The RNA data were log-normalized within every cell, and the ADT data were central-log-ratio-normalized across all cells. D. Violin plots comparing the expression levels of genes and proteins of interest in the two CD8+ T cell states. The unadjusted p-values were computed using Wilcoxon rank-sum tests. E. Violin plots comparing the expression levels of genes and proteins of interest in the two CD4+ T cell states. The unadjusted p-values were computed using Wilcoxon rank-sum tests.

**Supplementary Figure 3.**
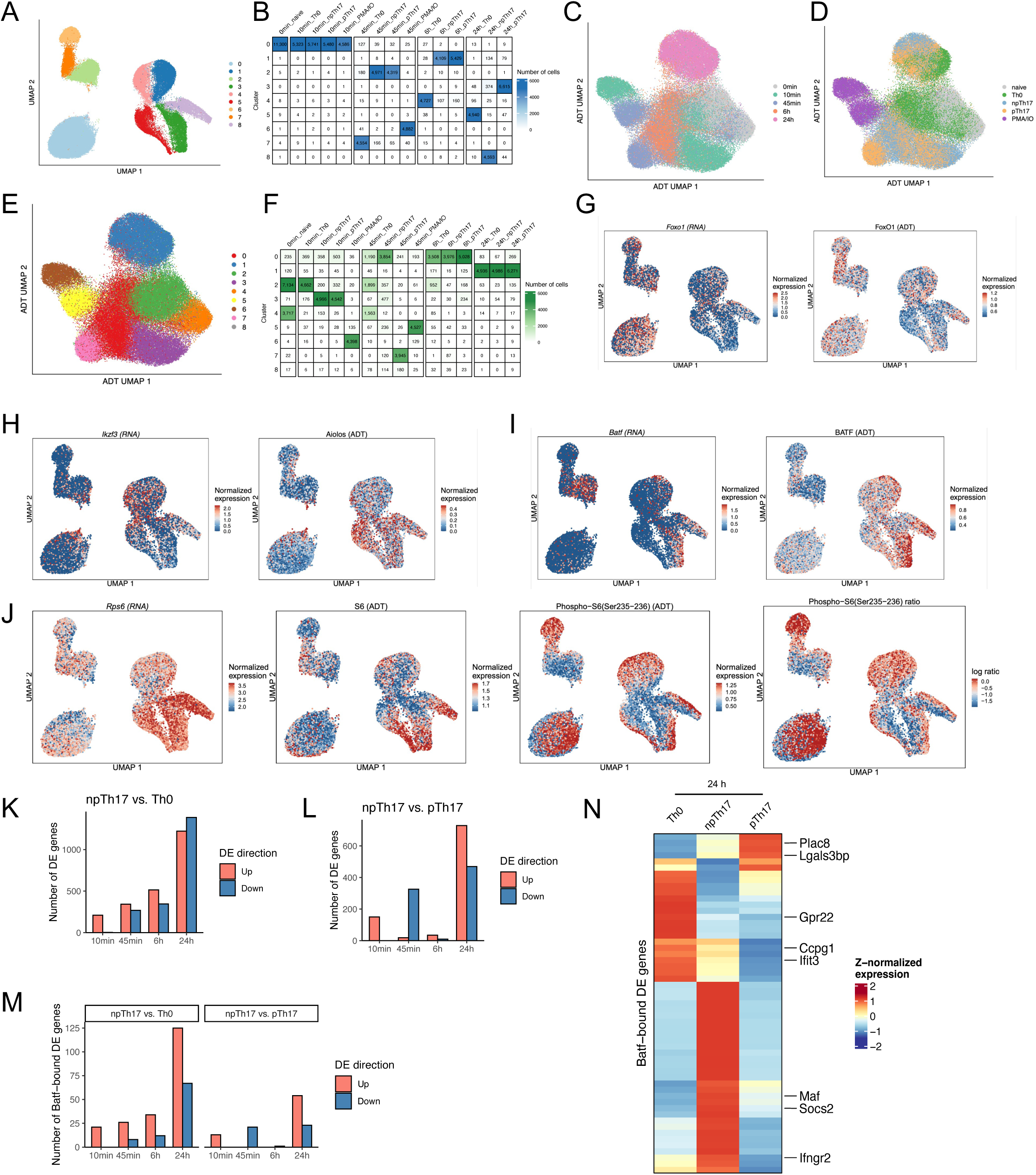
A. UMAP plot showing clusters identified based on RNA data. UMAP coordinates are the same as Figure 3B. Clustering was based on RNA data only. B. Number of cells in each cluster from different time points and stimulation conditions. Clusters are the same as Supplementary Figure 3A. C. UMAP plot showing the cells of different time points. UMAP coordinates were computed using protein (ADT) data only. D. UMAP plot showing the cells of different time points. The UMAP coordinates are the same as Supplementary Figure 3C. E. UMAP plot showing clusters identified based on RNA data. UMAP coordinates are the same as Supplementary Figure 3C. Clustering was based on protein data only. F. Number of cells in each cluster from different time points and stimulation conditions. Clusters are the same as Supplementary Figure 3E. G. UMAP plots depicting the expression of Foxo1 RNA and FoxO1 protein. H. UMAP plots depicting the expression of Ikzf3 RNA and Aiolos protein. I. UMAP plots depicting the expression of Batf RNA and BATF protein. J. UMAP plots depicting the expression of Rps6 gene (RNA), total S6 protein, and phospho-S6 (Ser235-236) protein, and the ratio of phospho-S6 (Ser235-236) to total S6 protein. K. Number of up- and down-regulated genes in npTh17 vs. Th0 at every time point. Differential expression test was performed using Wilcoxon rank-sum test, and threshold were set to FDR < 0.05 and fold change > 1.5. L. Number of up- and down-regulated genes in npTh17 vs. pTh17 at every time point. Differential expression test was performed using Wilcoxon rank-sum test, and threshold were set to FDR < 0.05 and fold change > 1.5. M. Number of Batf-bound genes that are significantly up- or down-regulated in npTh17 compared to Th0 (top) or pTh17 (bottom) at every time point. N. Expression of Batf direct targets of interest at the 24-hour time point. Batf direct targets of interest were defined as the Batf-bound genes that are both differentially expressed in npTh17 vs. Th0 at least one time point and are differentially expressed in npTh17 vs. pTh17 at least one time point. The color indicates the z-normalized average expression.

**Supplementary Figure 4.**
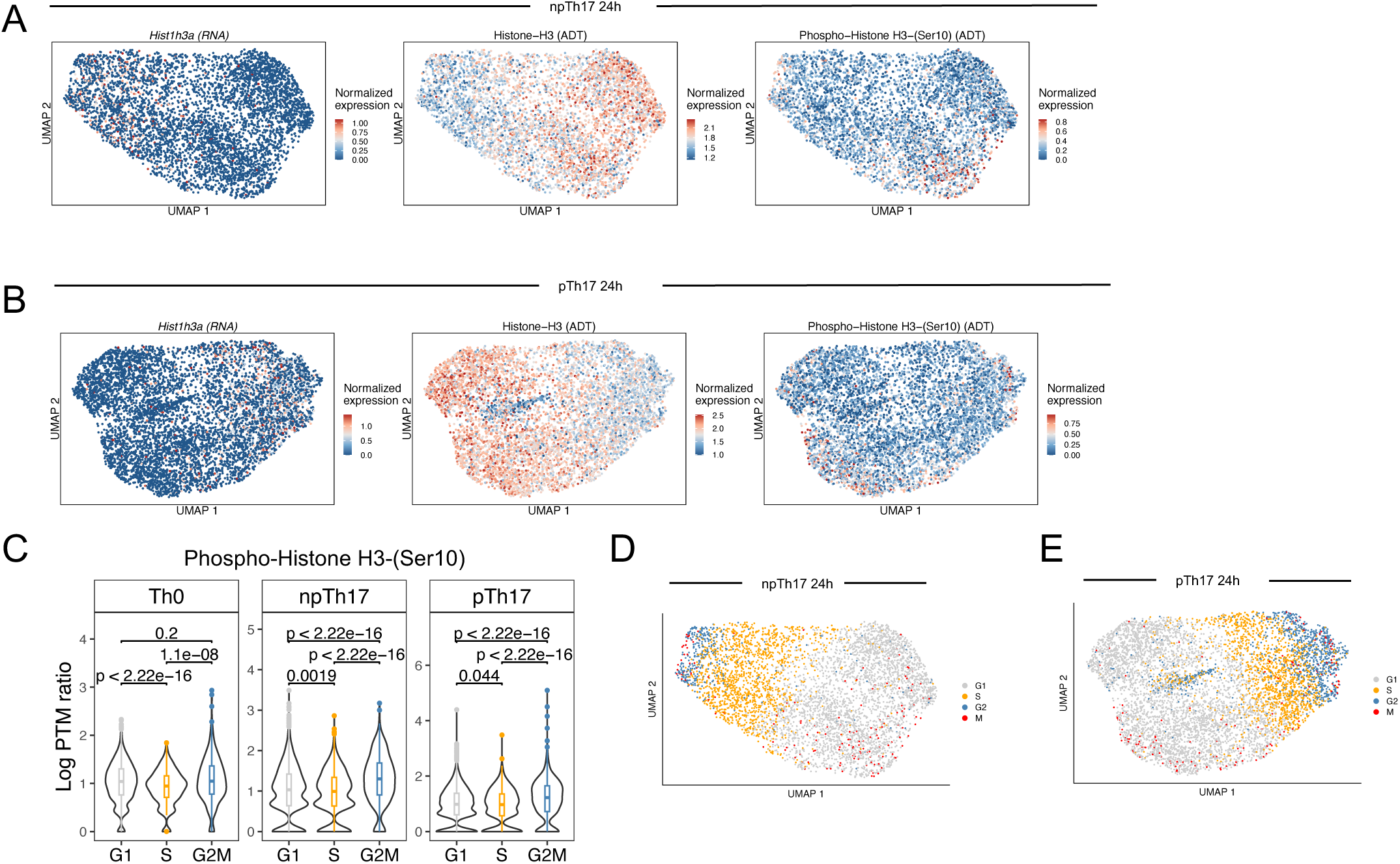
B. UMAP plots showing the expression of Hist1h3a gene (RNA), total Histone H3 protein, and pH3-Ser10 protein in npTh17 at 24 hours. C. UMAP plots showing the expression of Hist1h3a gene (RNA), total Histone H3 protein, and pH3-Ser10 protein in pTh17 at 24 hours. D. Distribution of pH3-Ser10 to total H3 protein ratios in cells at different cell cycle phases at 24 hours. E. UMAP plot showing the new cell cycle phase annotation using additional information from pH3-Ser10 protein levels in npTh17 at 24 hours. F. UMAP plot showing the new cell cycle phase annotation using additional information from pH3-Ser10 protein levels in pTh17 at 24 hours.

## Supplementary Tables

|  | Live | InTraSeq<br>RNA-only | InTraSeq<br>RNA+ADT |
| --- | --- | --- | --- |
| Total number of reads | 54,449,067 | 54,458,815 | 54,457,004 |
| Mean reads per cell | 7,778 | 7,780 | 7,780 |
| Median reads per cell | 5,126 | 5,547 | 5,867 |
| Mean UMIs per cell | 5,305 | 4,570 | 4,046 |
| Median UMIs per cell | 3,489 | 3,256 | 3,070 |
| Total number of genes | 26,647 | 27,300 | 27,171 |
| Mean genes per cell | 2,151 | 2,033 | 1,899 |
| Median genes per cell | 1,748 | 1,703 | 1,653 |

|  | Estimated.<br>Number.of.<br>Cells | Mean.Reads.<br>per.Cell | Median.Genes<br>per.Cell |
| --- | --- | --- | --- |
| 10min_npth17_gex | 7931 | 26309 | 1799 |
| 10min_PMA_IO_gex | 6213 | 18410 | 1552 |
| 10min_pTh17_gex | 7482 | 10992 | 1314 |
| 10min_Th0_gex | 7265 | 8739 | 1163 |
| 24h_npTh17_gex | 6871 | 15107 | 1921 |
| 24h_pTh17_gex | 9039 | 17998 | 1845 |
| 24h_Th0_gex | 6884 | 16658 | 1713 |
| 45min_npTh17_gex | 6825 | 10622 | 1134 |
| 45min_PMA_IO_gex | 6547 | 11803 | 1088 |
| 45min_pTh17_gex | 5716 | 15649 | 1305 |
| 45min_Th0_gex | 6499 | 11443 | 1145 |
| 6h_npTh17_gex | 5555 | 17037 | 1740 |
| 6h_pTh17_gex | 7351 | 13625 | 1481 |
| 6h_Th0_gex | 6406 | 12498 | 1456 |
| naive1_gex | 7186 | 8979 | 1086 |
| naive2_gex | 8359 | 11434 | 1304 |

